# Temporal Gut Microbial Changes Predict Recurrent *Clostridium difficile* in Patients with and without Ulcerative Colitis

**DOI:** 10.1101/632778

**Authors:** Allen A. Lee, Krishna Rao, Julajak Limsrivilai, Merritt Gillilland, Benjamin Malamet, Emily Briggs, Vincent B. Young, Peter DR Higgins

## Abstract

**Background:** Ulcerative colitis (UC) carries an increased risk of primary and recurrent *Clostridium difficile* infection (rCDI) and CDI is associated with UC flares. We hypothesized that specific fecal microbial changes associate with UC flare and rCDI.

**Methods:** We conducted a prospective observational cohort study of 57 patients with UC and CDI, CDI only, and UC flare only. Stool samples were collected at baseline, at the end of antibiotic therapy, and after reconstitution for 16S rRNA sequencing. The primary outcomes were recurrent UC flare and rCDI. Logistic regression and Lasso models were constructed for analysis.

**Results:** There were 21 (45.7%) patients with rCDI, while 11 (34.4%) developed UC flare. Patients with rCDI demonstrated significant inter-individual (*P*=.008) and intra-individual differences (*P*=.004 relative to baseline samples) in community structure by Jensen-Shannon distance (JSD) compared with non-rCDI. Two cross-validated models identified by Lasso regression predicted risk of rCDI: a baseline model with female gender, hospitalization for UC in the past year, increased Ruminococcaceae and Verrucomicrobia, and decreased Eubacteriaceae, Enterobacteriaceae, Lachnospiraceae, and Veillonellaceae (AuROC=0.94); and a model 14 days after completion of antibiotics with female gender, increased Shannon diversity, Ruminococcaceae and Enterobacteriaceae, and decreased community richness and Faecalibacterium (AuROC=0.9). Adding JSD between baseline and post-treatment samples to the latter model improved fit (AuROC=0.94). A baseline model including UC hospitalization in the past year and increased Bacteroidetes showed good fit characteristics for predicting increased risk of UC flare (AuROC=0.88).

**Conclusion:** Fecal microbial features at baseline and following therapy predict rCDI risk in patients with and without UC. These results may help risk stratify patients to guide management.

## INTRODUCTION

Although the pathogenesis of ulcerative colitis (UC) is incompletely understood, accumulating evidence suggests that disruptions of gut microbial structure and function contribute to inflammatory bowel disease (IBD). Introduction of gut microbiota taken from IBD patients and introduced into gnotobiotic mouse models of IBD can elicit proinflammatory immune responses similar to those seen in IBD patients.^1^ Moreover, intestinal inflammation may induce further derangements in the gut microbial community.^2, 3^ Thus, IBD may be seen as a deleterious feedback cycle of disruptions to the gut microbiota and alterations in host immune responses.

Patients with IBD, particularly those with UC, are at higher risk for *Clostridium difficile* (recently reclassified as Clostridiodes difficile) infection (CDI).^4–6^ IBD patients have multiple risk factors for CDI, including frequent antibiotic use, prior hospitalization, and/or a immunocompromised state, but CDI occurs at increased rates even in IBD patients without these traditional risk factors.^7^ It is possible that alterations in the gut microbial community may predispose IBD patients to CDI. Murine models suggest that shifts in microbial ecology are associated with susceptibility to experimental CDI.^8^ Notably, these changes in microbial dynamics in CDI are similar to those consistently observed in IBD. IBD patients with CDI also carry a 2-fold increased risk of hospitalization for subsequent exacerbation of IBD, increased risk for colectomy, and almost 5-fold increased risk for mortality compared to those without CDI.^5, 9–11^ IBD patients are also significantly more likely to have recurrent CDI (rCDI) compared with non-IBD controls.^12^

It is not clear how the microbiome disruptions seen in CDI and IBD relate to each other and/or interact. Thus, we aimed to characterize the fecal microbiota in patients with UC ± CDI longitudinally and investigate possible relationships to rCDI and recurrent UC flare. We hypothesized that poor reconstitution of the gut microbiome at the End of Antibiotics + 14 days (EOA+14) would be associated with rCDI and/or subsequent UC flare.

## METHODS

### Study Design

We conducted a prospective, observational cohort study at the University of Michigan. We recruited subjects from the following three groups: symptomatic patients with UC who also tested positive for CDI (cohort 1); non-IBD patients with symptomatic CDI (cohort 2); and patients with UC flare without CDI (cohort 3) (**Supplementary Figure 1**).

All subjects were 18 years or older and provided informed consent prior to enrollment in the study. Subjects were excluded from the study if they had presence of an ostomy or previous history of colectomy. Subjects in cohorts 1 and 3 had prior clinical, endoscopic and histologic diagnosis of UC while subjects in cohort 2 had no documented history of UC or autoimmune disease.^13^ The study was approved by the institutional review board at the University of Michigan (see Supplemental Methods for full details).

The primary outcomes were subsequent UC flare and rCDI. Secondary aims included identifying any microbial features at baseline that may discriminate between patients with CDI compared with UC. Subjects were contacted and their medical records were reviewed every 60 days for up to 180 days after enrollment to determine the recurrence of UC flare and rCDI. Due to the small sample sizes in each cohort, patients in cohorts 1 and 2 were analyzed collectively to determine the rate of rCDI while adjusting for UC status. Patients in cohort 1 were also followed to determine the rate of UC flare. As patients in cohort 3 (UC flare only) were experiencing an exacerbation of their UC on enrollment, it was not possible to differentiate between an on-going vs. recurrent UC flare. As a result, patients in cohort 3 were excluded from meeting the primary endpoint of UC flare but were used to adjust models for UC status. Patients in cohort 3 were also analyzed to determine potential differences in microbial variables between UC and CDI at baseline.

UC flare was defined by onset of typical symptoms occurring after enrollment in the study along with 6-point Mayo score > 2.5 and a fecal calprotectin > 150 in the absence of CDI.^14^ CDI was diagnosed by presence of diarrhea (≥ 3 unformed stools in a 24-hour period) and a positive stool test for toxigenic *C. difficile* (positive testing for both the glutamate dehydrogenase [GDH] antigen and TcdA/TcdB by EIA [C. Diff Quik Chek Complete®, Alere, Waltham, MA], or real-time PCR for the *tcdB* gene performed when GDH/toxin results were discordant [Simplexa™ *C. difficile* Universal Direct, Diasorin Molecular LLC, Cypress, CA]). Initial diagnostic tests were performed by the University of Michigan clinical microbiology laboratory. rCDI was defined by recurrence of symptoms at least 14 days after initial treatment of CDI and positive *C. difficile* testing.^15^

### Stool Collection

For cohorts 1 and 2, stool samples were collected from all subjects at baseline (day 0) prior to initiation of antibiotics for CDI and/or medical therapy for UC flare (**Supplemental Figure 2**). Stool samples were also collected at the end of antibiotics (EOA, approximately day 14) and EOA plus 14 days (approximately day 30). For cohort 3, stool samples were collected at baseline and at day 30 after clinical remission was achieved.

### 16S Sequence Analysis

DNA was extracted and libraries were prepared by the University of Michigan Host Microbiome core, the 16S rRNA genes were sequenced, and the *mothur* computational pipeline^16^ was deployed for processing sequence data as previously described (see Supplemental Methods).^17^ Following this, the following microbiome metrics were generated: Shannon diversity, Jensen-Shannon distance (JSD) relative to baseline and subsequent samples, community type using unsupervised partition around medioid clustering based on JSD,^18^ relative abundance of individual operational taxonomic units (OTUs), and other variables based on taxonomic class. One such constructed metric is the microbiome health index (MHI), defined as the proportion of Bacteroidia and Clostridia compared to the proportion of Gammaproteobacteria and Bacilli.^19^

### Statistical Analysis

Continuous data were reported as mean and standard deviation (SD) if normally distributed, and as median and range if not normally distributed. Categorical variables were reported as frequencies and percentages. Continuous data were compared using one-way ANOVA if data were parametric or Kruskal-Wallis test if non-parametric. Comparisons of proportions were performed using Fisher’s exact test. All data were analyzed using R version 3.5.2 (R Foundation for Statistical Computing, Vienna, Austria). A two-tailed P-value < 0.05 was considered significant for all analyses.

Unadjusted and adjusted logistic regression analyses were performed to identify clinical and microbial variables at baseline (time point 1) and at time point 3 (EOA plus 14 days) which were associated with subsequent UC flare and rCDI. Corrections for multiple comparisons were performed using the Benjamini-Hochberg method.^20^ Only OTUs that were present in at least 10% of samples were included in the analysis. The overall structure of microbial communities among our primary outcomes was compared using redundancy analysis, an ordination technique, followed by a permutational, multivariate ANOVA (PERMANOVA) for significance testing, as implemented by the R package *vegan* version 2.5-4.^21^

Two predictive models using different techniques and aims were performed. The first method included clinical and microbiota variables with P-values < .20 for the association with either rCDI or subsequent UC flare based on logistic regression results. A backward stepwise regression method was used to select predictors in the final multivariable model, and interactions among the variables in the final model were assessed. This modeling strategy helped quantify the magnitude, strength, and statistical significance of individual predictors while accounting for confounding. However, it is not ideal for avoiding overfitting and maximizing generalizability of models.

Due to the large number of possible predictor variables, and in order to generate models that minimized overfitting, a second approach using Lasso (least absolute shrinkage and selection operator) regression with cross validation was also employed. Models were built in a stepwise regression fashion, and the optimal model was automatically selected using a 3-fold cross-validation that minimized the penalty term (i.e. λ), as implemented in the *glmnet* package version 2.0-16.^22^ Since cross-validation includes a component of randomness, this stepwise modeling strategy was simulated 1,000 times and those variables that appeared most frequently were selected for inclusion in the final model. The area under the receiver operator characteristic curve (AuROC) was calculated for each model using the R package pROC version 1.14.0.^23^

To assess longitudinal associations between clinical and microbial variables of interest with the primary outcomes, generalized estimating equations (GEE) with an exchangeable or auto-regressive correlation structure, generalized linear mixed-effects models (GLMER) and generalized additive models (GAM) were utilized using the R packages *geepack* version 1.2-1,^24^ *lme4* version 1.1-21,^25^ and *mgcv* version 1.8-28,^26^ respectively.

## RESULTS

### Baseline Clinical Variables

A total of 57 subjects were enrolled in this study (32 with UC/CDI, 14 with CDI only, and 11 with UC only). Patients with CDI only were older compared to those with UC/CDI and UC only (*P*=.001) (Table 1). Patients with UC only were significantly more likely to receive steroids as their initial treatment for UC flare compared to those subjects with UC and CDI (90.9% vs. 31.2%, *P*=.001). Patients with UC only were also less likely to have received antibiotics in the past year compared to patients with UC and CDI or patients with CDI only (*P*=.006). There were no other clinical variables at baseline that differed significantly between the three groups. A total of 21 subjects (45.7%) met the primary endpoint for rCDI while 11 subjects (34.4%) developed a subsequent UC flare.

**Table 1.**
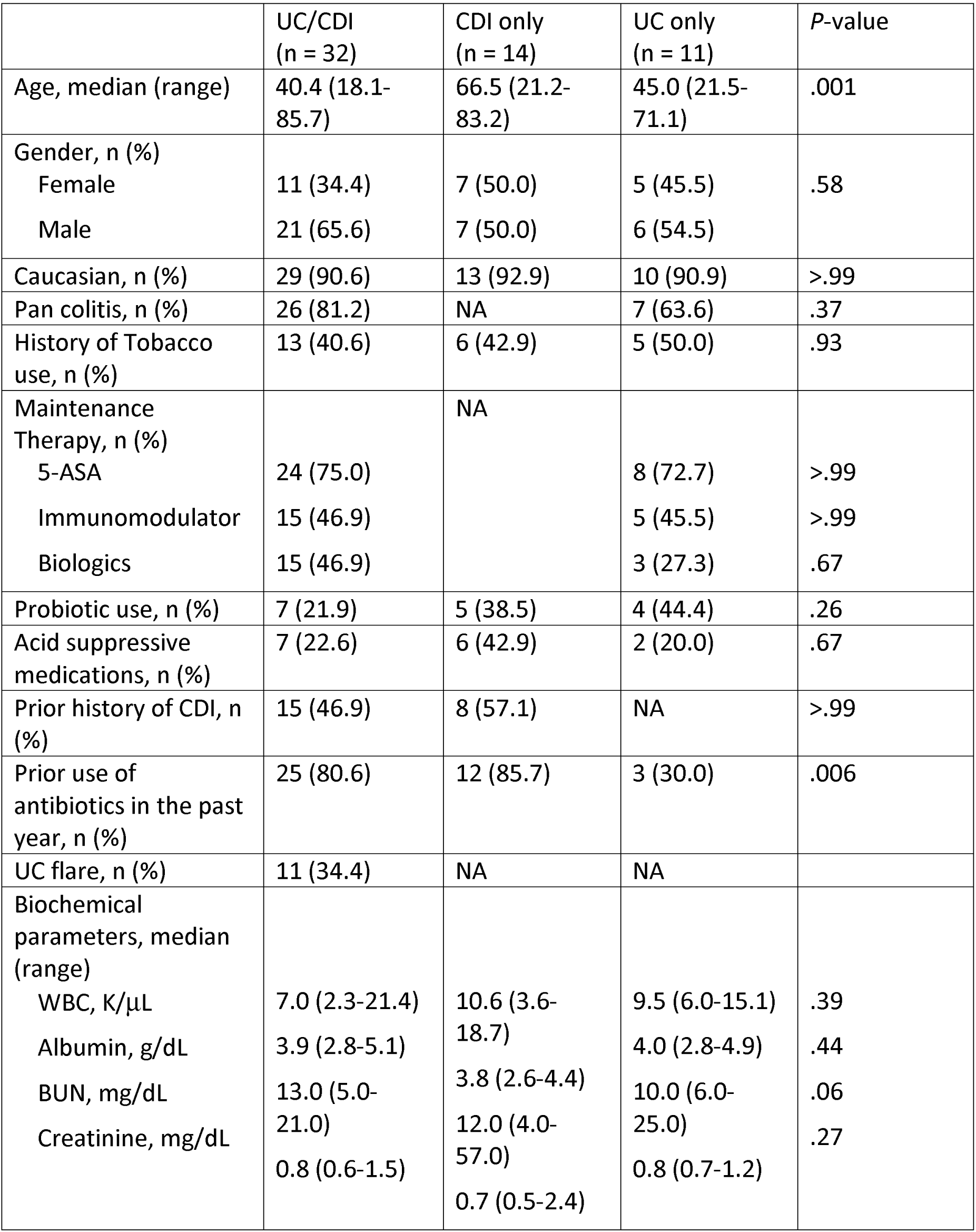
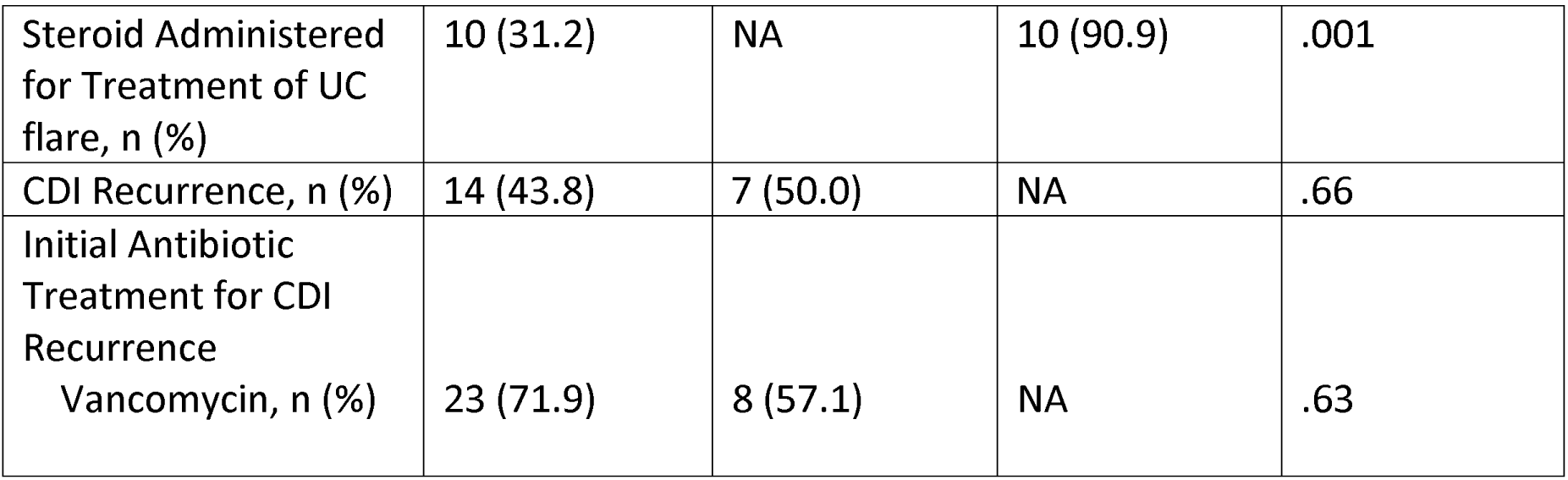
Baseline Demographic Variables for Patients with Ulcerative Colitis (UC) with or without *Clostridium difficile* Infection (CDI).

|  | UC/CDI<br>(n = 32) | CDI only<br>(n = 14) | UC only<br>(n = 11) | P-value |
| --- | --- | --- | --- | --- |
| Age, median (range) | 40.4 (18.1-85.7) | 66.5 (21.2-83.2) | 45.0 (21.5-71.1) | .001 |
| Gender, n (%) |  |  |  |  |
| Female | 11 (34.4) | 7 (50.0) | 5 (45.5) | .58 |
| Male | 21 (65.6) | 7 (50.0) | 6 (54.5) |  |
| Caucasian, n (%) | 29 (90.6) | 13 (92.9) | 10 (90.9) | >.99 |
| Pan colitis, n (%) | 26 (81.2) | NA | 7 (63.6) | .37 |
| History of Tobacco use, n (%) | 13 (40.6) | 6 (42.9) | 5 (50.0) | .93 |
| Maintenance Therapy, n (%) |  | NA |  |  |
| 5-ASA | 24 (75.0) |  | 8 (72.7) | >.99 |
| Immunomodulator | 15 (46.9) |  | 5 (45.5) | >.99 |
| Biologics | 15 (46.9) |  | 3 (27.3) | .67 |
| Probiotic use, n (%) | 7 (21.9) | 5 (38.5) | 4 (44.4) | .26 |
| Acid suppressive medications, n (%) | 7 (22.6) | 6 (42.9) | 2 (20.0) | .67 |
| Prior history of CDI, n (%) | 15 (46.9) | 8 (57.1) | NA | >.99 |
| Prior use of antibiotics in the past year, n (%) | 25 (80.6) | 12 (85.7) | 3 (30.0) | .006 |
| UC flare, n (%) | 11 (34.4) | NA | NA |  |
| Biochemical parameters, median (range) |  |  |  |  |
| WBC, K/μL | 7.0 (2.3-21.4) | 10.6 (3.6-18.7) | 9.5 (6.0-15.1) | .39 |
| Albumin, g/dL | 3.9 (2.8-5.1) | 3.8 (2.6-4.4) | 4.0 (2.8-4.9) | .44 |
| BUN, mg/dL | 13.0 (5.0-21.0) | 12.0 (4.0-57.0) | 10.0 (6.0-25.0) | .06 |
| Creatinine, mg/dL | 0.8 (0.6-1.5) | 0.7 (0.5-2.4) | 0.8 (0.7-1.2) | .27 |
| Steroid Administered for Treatment of UC flare, n (%) | 10 (31.2) | NA | 10 (90.9) | .001 |
| CDI Recurrence, n (%) | 14 (43.8) | 7 (50.0) | NA | .66 |
| Initial Antibiotic Treatment for CDI Recurrence<br>Vancomycin, n (%) | 23 (71.9) | 8 (57.1) | NA | .63 |

### Patients with CDI Show Reduced Microbial Diversity and Richness at Baseline Compared with UC

Microbial features at baseline were compared in patients with UC and CDI. Patients with CDI showed decreased Shannon diversity (P<.05) and community richness (*P*=.002) at baseline compared with UC even after controlling for UC status (Supplemental Table 1). There were also several OTUs, including several Lachnospiraceae genera, that were depleted at baseline in patients with CDI compared with UC.

### Patients with rCDI Exhibited a Distinct Community Structure Compared with Non-rCDI

We next performed redundancy analysis to explore differences in microbial communities across populations with rCDI and UC flare as variables (cohorts 1 and 2). There were significant differences in the baseline community structure between patients who subsequently developed rCDI vs. non-rCDI (*P*=.008 by PERMANOVA), Figure 1A). No significant differences were seen between patients with subsequent UC flare vs. without a recurrent UC flare (*P*=.44) (Figure 1B).

**Figure 1.**
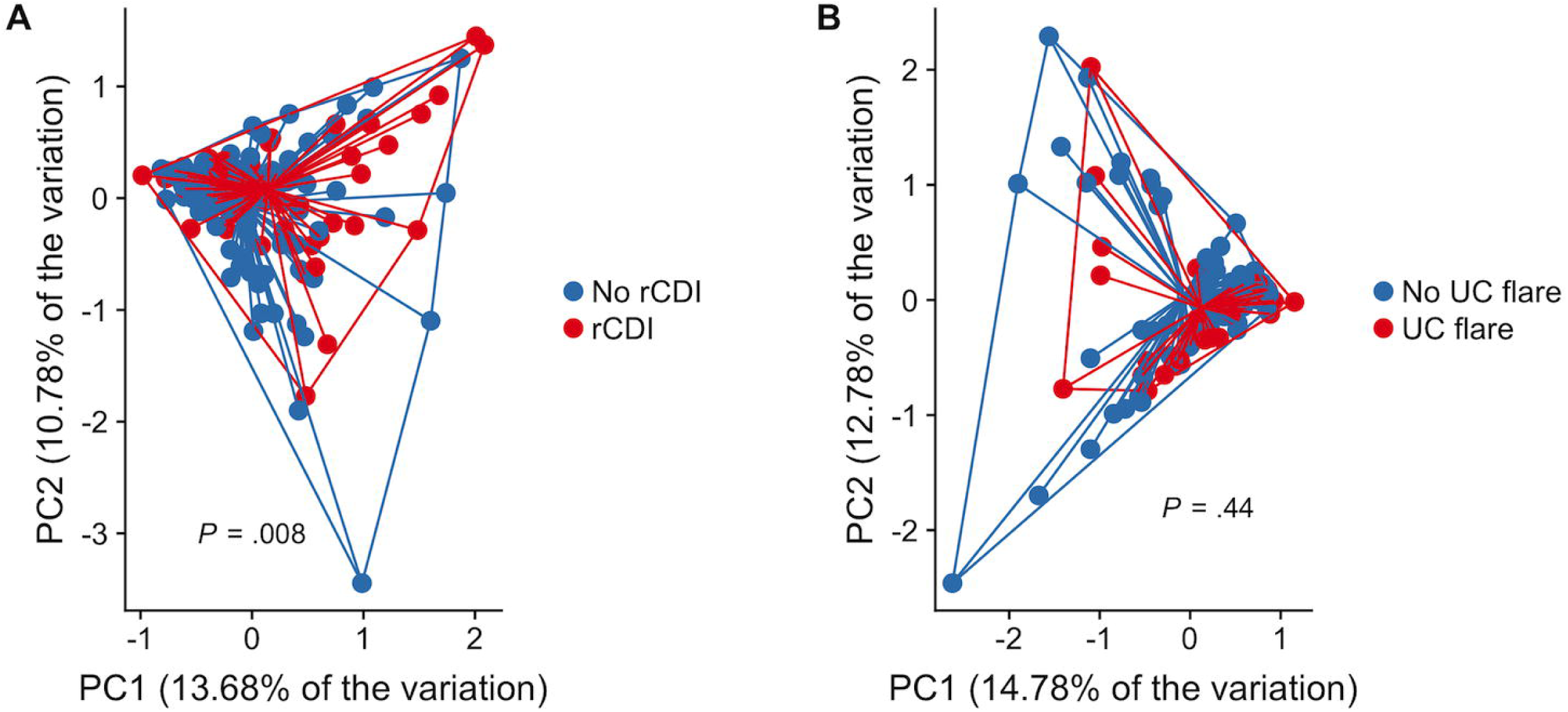
Patients with Recurrent *Clostridium difficile* Infection (rCDI) Show Broad Differences in Microbial Community Structure Compared with non-rCDI. (A) Principal coordinates analysis (PCoA) based on redundancy analysis demonstrated significant differences in the community structure between patients who subsequently developed rCDI (red) compared with patients without rCDI (blue) (*P*=.008 by PERMANOVA). (B) PCoA plot showing no differences between community structures in patients who subsequently developed UC flare (red) vs. non-UC flare (blue) (*P*=.44).

### Patients with rCDI Showed Greater Intra-Individual Variability of the Fecal Microbiota Over Time

We then evaluated intra-individual community changes in cohorts 1 and 2 by Jensen-Shannon Distance (JSD, ranging from 0, indicating complete similarity relative to baseline samples, to 1 or complete dissimilarity with baseline samples) over time. At the end of antibiotics (EOA), there were no differences in JSD between patients who developed rCDI and those who did not develop rCDI (*P*=.41) (Figure 2A). However, 14 days after completion of antibiotics, those patients who subsequently developed rCDI demonstrated greater dissimilarity to baseline samples in community structure compared with non-rCDI patients (*P*=.004) (Figure 2B). There were no differences in JSD at EOA (*P*=.75) or at EOA plus 14 days (*P*=.22) in patients with and without subsequent UC flare.

**Figure 2.**
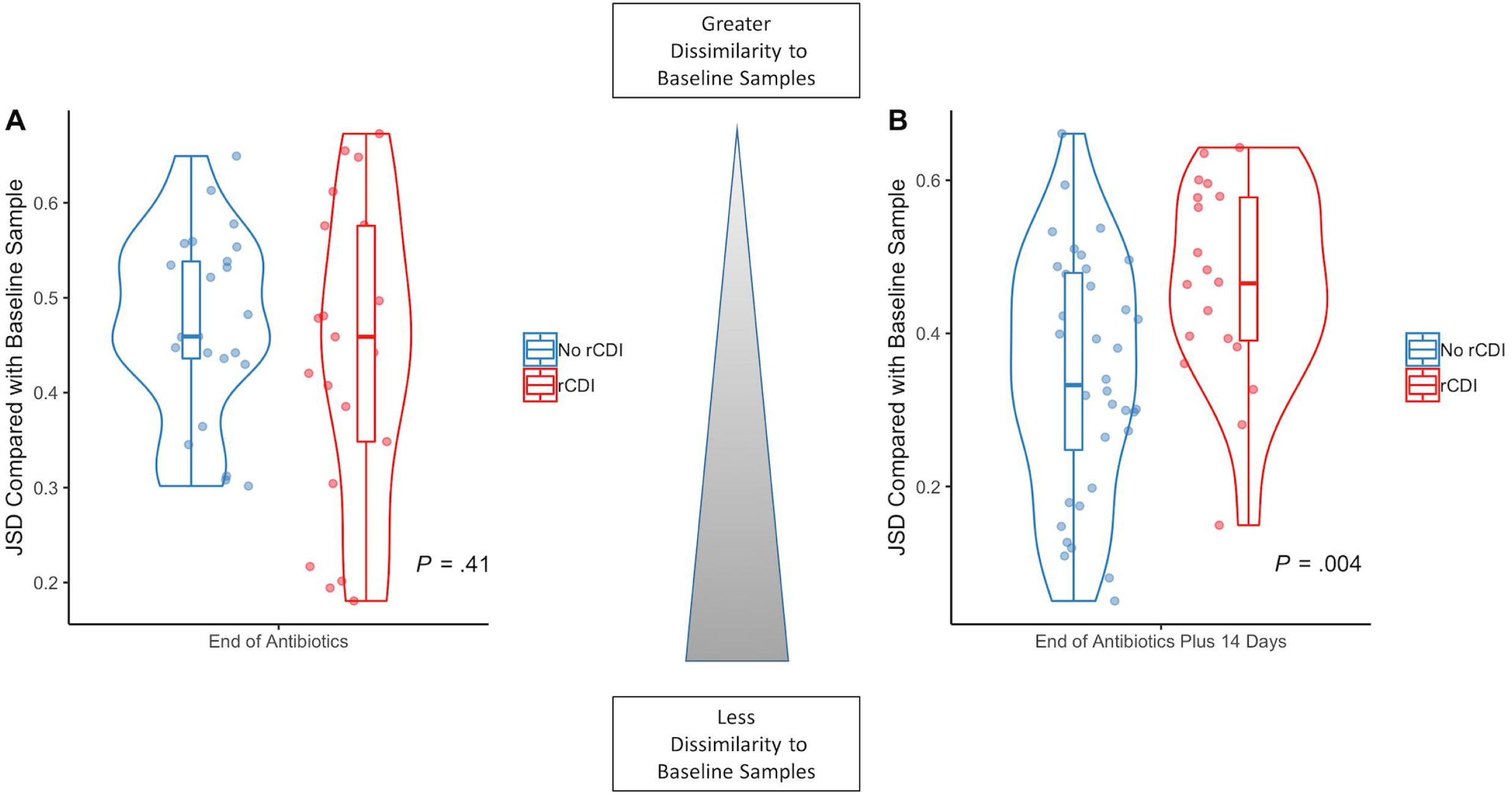
Patients with Recurrent *Clostridium difficile* Infection (rCDI) Show Significant Perturbations of the Fecal Microbiota Longitudinally Compared with Non-rCDI Patients. Jensen-Shannon distance (JSD), a measure of intra-individual microbial community dissimilarity compared to baseline samples, was calculated over time. (A) At the end of antibiotics, there were no differences in JSD between patients who subsequently developed rCDI (red) and non-rCDI (blue) (*P*=.41). (B) However, 14 days after completion of antibiotics, patients who subsequently developed rCDI (red) demonstrated a significantly elevated JSD (greater community dissimilarity relative to baseline samples) compared with non-rCDI patients (blue) (*P*=.004). These results suggest that patients with on-going perturbations of the fecal microbiota 2 weeks after completion of antibiotics were at higher risk for rCDI.

### Baseline Clinical and Microbial Variables Are Predictive for rCDI

#### Unadjusted Variables Associated with rCDI

We performed logistic regression models to determine baseline variables that were associated with risk for rCDI (cohorts 1 and 2) while controlling for UC status (cohorts 1 and 3) (Supplemental Table 2). Female gender was associated with an increased risk for rCDI (OR=2.5, *P*=.05). In terms of microbial variables, an increase in the relative abundance of Lachnospiraceae at baseline was protective against subsequent risk for rCDI (OR=0.52 for every 10% increase, *P*=.02). There were no other clinical or microbial variables at baseline that were associated with risk for rCDI.

#### Multivariable Model for rCDI

Using backward stepwise regression analyses, in our final model we identified five baseline variables that were significantly associated with rCDI even after adjusting for UC status (Table 2). Female gender was associated with an increased risk for rCDI (OR=16.2, *P*=.005). In contrast, increased OTU richness (OR=0.86 per every increase of 10 taxa, *P*=.02) as well as increased relative abundance of Enterobacteriaceae (OR=0.29 per every 10% increase, *P*=.004); Lachnospiraceae (OR=0.17 per every 10% increase, *P*=.002); and Veillonellaceae (OR=0.17 per every 10% increase, *P*=.05) were protective against rCDI (Figure 3A-D). We assessed for interactions among variables in the final model and none were found. This final model had excellent fit characteristics (AuROC=0.91) (Figure 3E).

**Figure 3.**
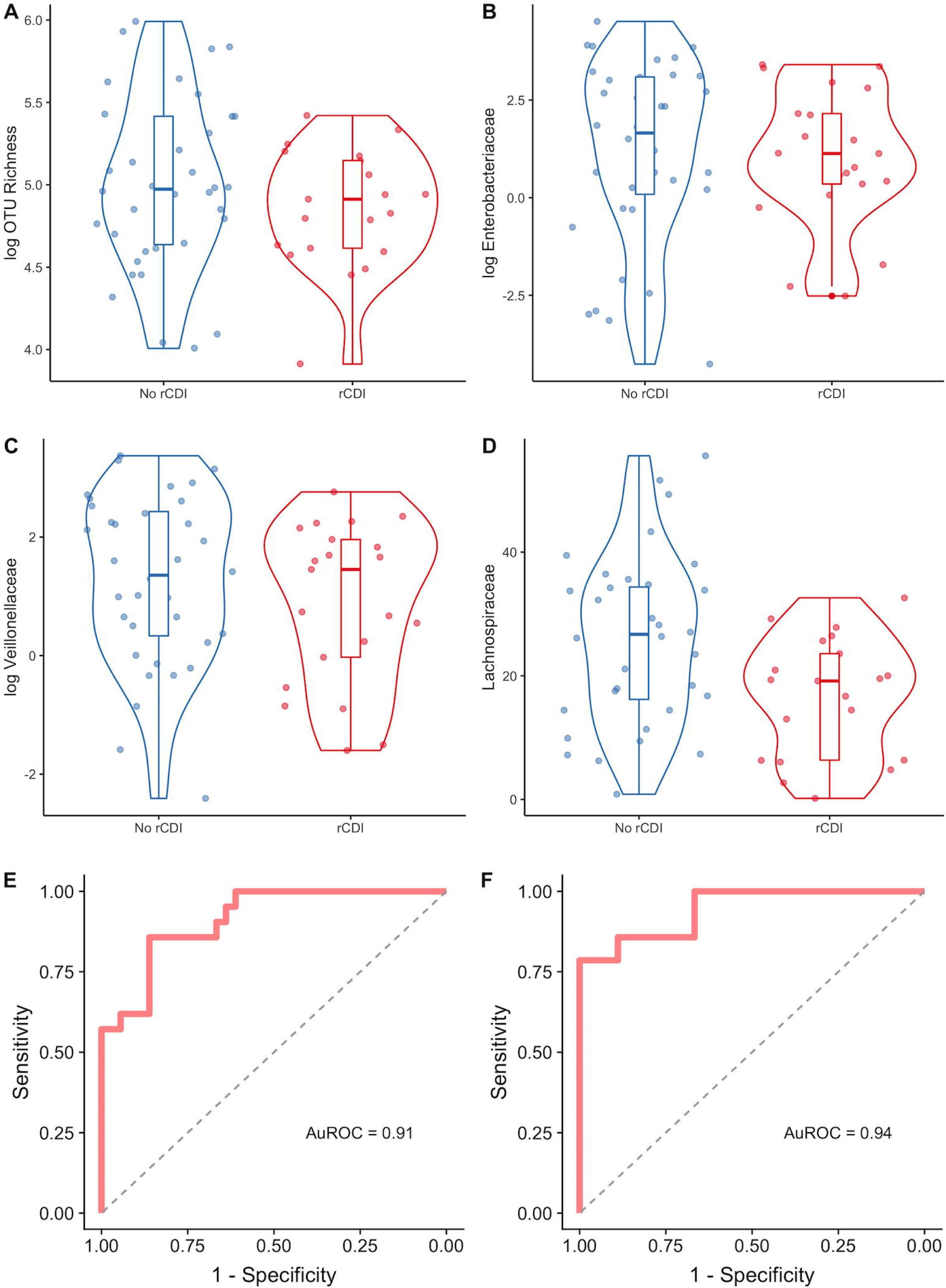
Identification of Microbial Variables at Baseline Associated with Increased or Decreased Risk for Recurrent *Clostridium difficile* Infection (rCDI). Three bacterial taxa and community richness were identified by backward stepwise regression as being associated with subsequent rCDI. The relative abundances for these three taxa as well as community richness are shown grouped by whether or not rCDI occurred. Data are shown as violin plots with accompanying boxplot and whiskers indicating median, interquartile range (IQR) and 1.5 x IQR of the median for the following variables: (A) OTU richness, (B) Enterobacteriaceae, (C) Lachnospiraceae, and (D) Veillonellaceae. Note that OTU richness, Enterobacteriaceae, and Veillonellaceae are shown in logarithmic scale. (E) Receiver Operating Characteristic (ROC) Curve for this model identified by backward stepwise regression is shown (AuROC=0.91). (F) ROC for model selected by Lasso regression, which included variables for female gender, hospitalization for UC in the previous year, and increased relative abundances of Ruminococcaceae and Verrucomicrobia, and decreased Eubacteriaceae, Enterobacteriaceae, Veillonellaceae, and Lachnospiraceae (AuROC=0.94).

**Table 2.** Multivariable Logistic Regression Models of Clinical and Microbial Variables at Baseline Associated with Recurrent *Clostridium difficile* Infection (rCDI).

|  | OR | 95% CI | P-value |
| --- | --- | --- | --- |
| ***Clinical Variables Associated with Increased Risk for rCDI*** |  |  |  |
| Female gender | 16.2 | 2.77 – 155.62 | .005 |
| ***Microbial Variables Protective Against rCDI*** |  |  |  |
| OTU Richness (per every increase of 10 taxa) | 0.86 | 0.73 – 0.96 | .02 |
| Enterobacteriaceae (per every 10% increase) | 0.29 | 0.09 – 0.56 | .004 |
| Lachnospiraceae (per every 10% increase) | 0.17 | 0.04 – 0.44 | .002 |
| Veillonellaceae (per every 10% increase) | 0.17 | 0.02 – 0.75 | .05 |

Effect sizes are presented after adjustment for UC status.

Model selection using 3-fold cross-validated Lasso regression to minimize λ with 1000 simulations identified two clinical and six taxonomic variables at baseline that were associated with rCDI in the largest number of simulations (Supplemental Table 3). Increased risk for rCDI was associated with hospitalization for UC in the past year and female gender, as well as increased relative abundances of Verrucomicrobia and Ruminococcaceae. Conversely, an increase in Eubacteriaceae, Lachnospiraceae, Veillonellaceae, and Enterobacteriaceae at baseline were protective against subsequent risk for rCDI. When these variables were included in a Lasso regression model, this final model demonstrated excellent predictive capabilities for rCDI (AuROC=0.94) (Figure 3F).

### Baseline Clinical and Microbial Variables Associated with Increased Risk for UC Flare

#### Unadjusted Variables Associated with UC flare

Prior hospitalization for UC in the past year (OR=16.0, *P*=.003) as well as steroid use as initial treatment for UC exacerbation (OR=10.5, *P*=.008) were significantly associated with risk for UC flare (cohort 1) (Supplemental Table 2). An increase in the relative abundance of Bacteroidetes was associated with increased risk for UC flare (OR=2.06 for every 10% increase, *P*=.01) (**Supplemental Figure 3**).

#### Multivariable Modeling Associated with UC Flare

Using backward stepwise regression, hospitalization for UC in the previous year (OR=17.70, *P*=.008) and increased relative abundance of Bacteroidetes at baseline (OR=1.07, *P*=.03) were associated with increased risk for UC flare (AuROC=0.88).

### Longitudinal Models Utilizing Microbial Variables Predict Increased Risk for rCDI

We used microbial data from all timepoints in cohorts 1 and 2 to model rCDI by fitting generalized estimating equations (GEE) and generalized linear mixed models (GLMM) while adjusting for UC status (cohorts 1 and 3). However, models using both GEE and GLMM either did not converge, resulted in singular fits, or gave nonsensical results. This was likely related to excessive collinearity of variables, overdispersion/excessive variability over time, and/or nonlinearity in the shape of the data. In terms of the latter, since the data did not follow a linear pattern (**Supplemental Figure 4A-D**), we tried three different approaches to produce a coherent model. First, in order to more accurately model the parabolic nature of the dataset, a quadratic term was introduced but this still did not result in models that converged. Secondly, we also modeled our data at two different time points, including at time points 1 (baseline) and 3 (EOA plus 14 days) and separately at time points 2 (EOA) and 3 (EOA plus 14 days). However, modeling with only two time points did not produce a model that converged. Finally, we used generalized additive models (GAM), which introduce a penalized regression spline in order to model non-linear data. Here, we identified by GAM that female gender as well as decreasing OTU richness and increasing Ruminococcaceae longitudinally were associated with future risk for rCDI (AuROC=0.87).

### Clinical and Microbial Variables at Time Point 3 Also Predict Risk for rCDI

#### Unadjusted Variables Associated with rCDI

We next performed logistic regression modeling to determine whether microbial features at time point 3 (microbial reconstitution, EOA plus 14 days) were associated with rCDI (cohorts 1 and 2) while controlling for UC status (cohorts 1 and 3) (Supplemental Table 4). An increased relative abundance of Gammaproteobacteria (OR=1.63 per every 10% increase, *P*=.04), Enterobacteriaceae (OR=1.78 per every 10% increase, *P*=.03), and Jensen-Shannon Distance (JSD) at time point 3 relative to baseline (OR=1.80 per every 0.1 increase, *P*=.01) were associated with increased rCDI risk. Conversely, an increase in Shannon diversity (OR=0.34, *P*=.04) and OTU richness (OR=0.85 per every increase of 10 taxa, *P*=.008) as well as increased relative abundance of Ruminococcaceae (OR=0.40 per every 10% increase, *P*=.04) and Faecalibacterium (OR=0.10 per every 10% increase, *P*=.04) were all protective against future risk for rCDI. Additionally, at a lower level of significance, increased Bacteroidetes (OR=0.60 per every 10% increase, *P*=.05), Lachnospiraceae (OR=0.62 per every 10% increase, *P*=.06) and MHI (OR=0.78 per every 1% increase, *P*=.05) were also protective against rCDI.

#### Multivariable Model Selection for rCDI

Model selection by backward stepwise regression identified four clinical and microbial variables at time point 3 that were significantly associated with rCDI after adjusting for UC status (Table 3). Female gender (OR=5.69, *P*=.03) and increased relative abundance of Ruminococcaceae (OR=1.43 per every 1% increase, *P*=.03) were associated with increased risk for future rCDI. In contrast, an increase in OTU richness (OR=0.83 per every increase of 10 taxa, *P*=.03), and increased relative abundance of Faecalibacterium (OR=0.47 per every 1% increase, *P*=.02) were protective against rCDI (Figure 4A-C). This model showed good fit characteristics (AuROC=0.87) for predicting rCDI. When Jensen-Shannon Distance (JSD) at time point 3 relative to baseline was added to the model, it had a statistically significant effect by likelihood ratio testing (*P*=.007) and improved the model fit (AuROC=0.90).

**Figure 4.**
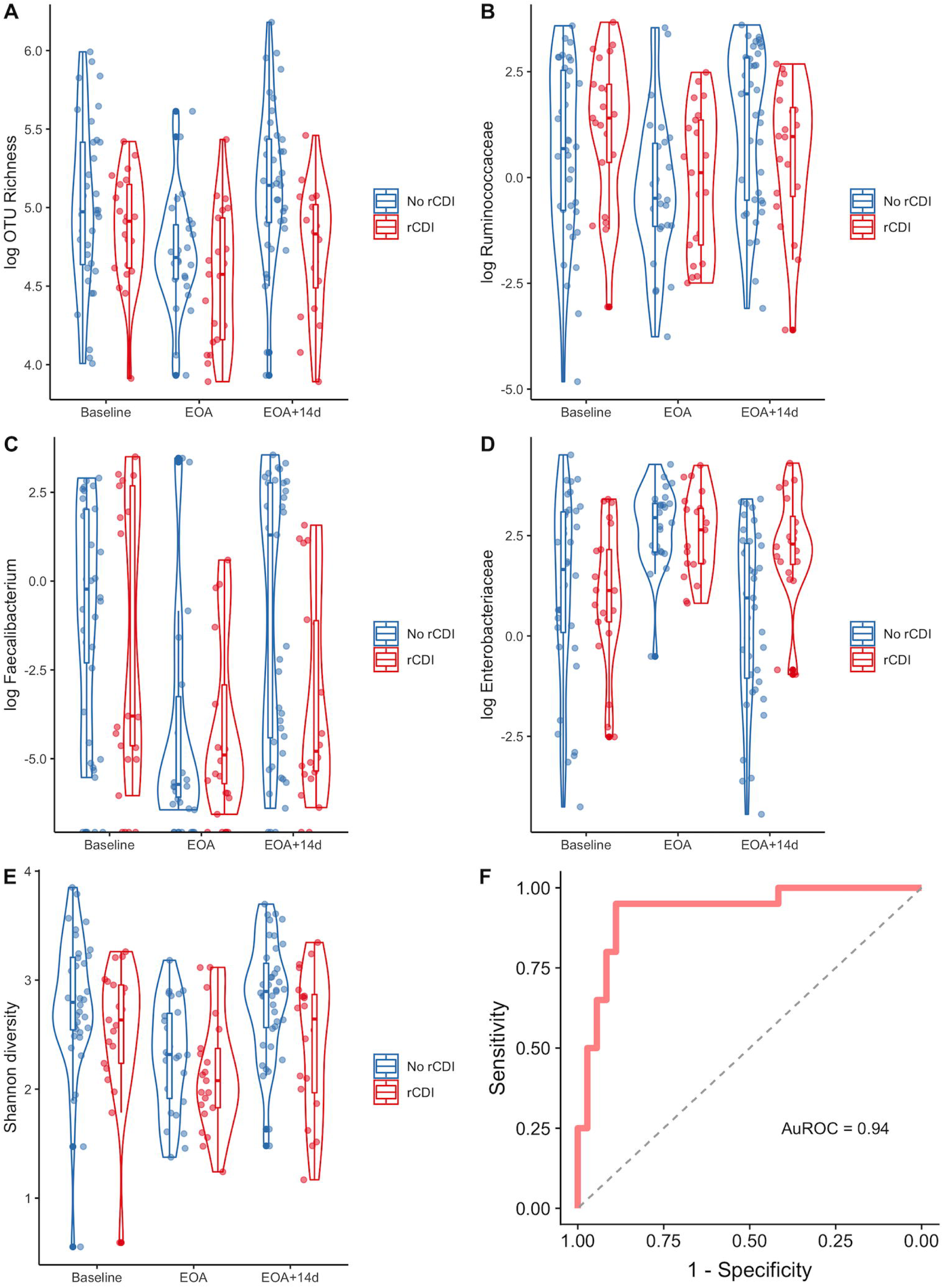
Fecal Microbial Characteristics at End of Antibiotics Plus 14 Days are Associated with Future Risk for Recurrent CDI. Several microbial variables 14 days after completion of antibiotics were identified as being associated with risk for future rCDI. The relative abundances for representative taxa as well as Shannon diversity and community richness are shown grouped by whether or not rCDI occurred. Data are shown as violin plots with accompanying boxplot and whiskers indicating median, interquartile range (IQR) and 1.5 x IQR of the median for the following variables: (A) OTU richness, (B) Ruminococcaceae, (C) Faecalibacterium, (D) Enterobacteriaceae, and (E) Shannon diversity over time, including at baseline, end of antibiotics (EOA), and end of antibiotics plus 14 days (EOA+14d). Note that OTU richness, Ruminococcaceae, Faecalibacterium, and Enterobacteriaceae are shown in logarithmic scale. (F) ROC curve displaying good fit characteristics of model identified by Lasso regression containing variables of female gender, increased Shannon diversity, increased relative abundances of Ruminococcaceae and Enterobacteriaceae, decreased OTU richness and Faecalibacterium, and Jensen Shannon Distance (relative to baseline) at end of antibiotics plus 14 days for risk of future rCDI (AuROC=0.94).

**Table 3.** Multivariable Logistic Regression Models of Clinical and Microbial Variables at Time Point 3 (14 days after Completion of Antibiotics) Associated with Recurrent *Clostridium difficile* Infection (rCDI).

|  | OR | 95% CI | P-value |
| --- | --- | --- | --- |
| ***Clinical and Microbial Variables Associated with Increased Risk for rCDI*** |  |  |  |
| Female gender | 5.69 | 1.36 – 29.71 | .03 |
| Ruminococcaceae (per every 1% increase) | 1.43 | 1.08 – 2.07 | .03 |
| Jensen Shannon Divergence (per every 0.1 increase) | 2.59 | 1.28 – 6.15 | .01 |
| ***Microbial Variables Protective Against Risk for rCDI*** |  |  |  |
| OTU Richness (per every increase of 10 taxa) | 0.83 | 0.66 – 0.96 | .03 |
| Faecalibacterium (per every 1% increase) | 0.47 | 0.23 – 0.75 | .02 |

Effect sizes are presented after adjustment for UC status.

Cross-validated Lasso regression simulations identified similar variables to the backward stepwise regression, including an increased relative abundance of Ruminococcaceae, decreased OTU richness, and decreased Faecalibacterium at time point 3 (EOA plus 14 days) as being associated with increased risk for rCDI (Supplemental Table 5). Lasso regression also identified additional variables associated with increased rCDI, including female gender, increased Enterobacteriaceae (Figure 4D), and increased Shannon diversity (Figure 4E). When these variables selected by the greatest number of simulations were included in a final Lasso regression model, this model showed excellent predictive capabilities for identifying rCDI (AuROC=0.90). When Jensen-Shannon Distance (JSD) at time point 3 relative to baseline was added, the model showed increased fit characteristics (AuROC=0.94) (Figure 4F).

### Clinical and Microbial Variables Did Not Associate with UC Flare at Time Point 3

We did not identify any microbial variables at time point 3 in cohort 1 that were associated with risk for UC flare. Similarly, model selection by Lasso regression did not identify any variables that were significantly associated with UC flare (data not shown).

## DISCUSSION

We have conducted a prospective, longitudinal cohort study examining whether gut microbial changes can predict future risk for rCDI and recurrent flare of UC in a cohort of patients with CDI, with and without UC. In this study, we have shown that specific microbial characteristics, either at baseline or upon reconstitution of the microbiome, can detect those patients at high risk for future episodes of rCDI and UC flare. Specifically, we demonstrated that (1) patients with CDI ± UC who subsequently develop rCDI possess a distinct microbial community structure compared with non-rCDI patients; (2) there are unique microbial characteristics at baseline which can identify patients at risk for rCDI with 94% accuracy; (3) those patients with on-going perturbations of their fecal microbiota two weeks after completion of antibiotics are at increased risk for rCDI; and (4) patients with history of hospitalization for UC in the previous year and increased Bacteroidetes at baseline had increased risk for subsequent UC flare.

Patients with UC and CDI represent a high-risk patient population with significant risk for rCDI and poor outcomes. Identifying those patients with higher risk for recurrence of disease would represent a major advance in the field. Our results have identified several key variables at baseline that are associated with rCDI even after adjusting for UC status. Specifically, female gender, hospitalization for UC in the prior year, increased Ruminococcaceae and Verrucomicrobia were associated with increased risk for rCDI while increased abundances of Eubacteriaceae, Enterobacteriaceae, Lachnospiraceae, and Veillonellaceae were protective against risk for future rCDI. This model demonstrated excellent discriminative ability with 94% accuracy to predict future occurrences of rCDI. Notably, these variables were identified in baseline samples, which suggests that high-risk patients can be identified at the time of CDI diagnosis. These patients might benefit from more aggressive therapy, such as prolonged antibiotic tapers, an antibiotic with lower risk of rCDI such as fidaxomicin,^27^ use of a monoclonal antibody to reduce risk of rCDI,^28^ fecal microbiota transplantation (FMT), or even a rationally designed probiotic (based on data from the current study) containing Eubacteriaceae, Lachnospiraceae, Enterobacteriaceae, and Veillonellaceae.

Female gender has previously been identified as an independent risk factor for CDI,^29^ which may be related to differences in sex hormone levels resulting in differences in gut microbial composition.^30^ Widespread changes in the gut microbiota, including lower diversity and decreased abundance of putatively beneficial bacteria, such as Lachnospiraceae, have been demonstrated in both UC and CDI.^31, 32^ Lachnospiraceae is a primary producer of butyrate, which is known to inhibit *C. difficile in vitro*^33^ and enhances intestinal epithelial barrier function.^34^ Furthermore, both *C. difficile* and Lachnospiraceae are taxonomically classified within the Clostridiales order. Thus, Lachnospiraceae may be protective against CDI by occupying a similar ecological niche within the gastrointestinal tract competing for similar resources as C. difficile. In addition, a probiotic containing 33 different bacterial species, including Lachnospiraceae, Ruminococcaceae, and Eubacteriaceae, was protective against rCDI for up to 6 months posttreatment.^35^

Although there is a paucity of data regarding association between microbial characteristics and rCDI, the available data show some differences from our results. Studies prior to FMT in CDI patients have demonstrated widespread changes in the gut microbiota, including decreased abundances of Lachnospiraceae and Ruminococcaceae as well as enrichment of Enterococcaceae and Veillonellaceae.^36^ Khanna, *et al.* reported increased abundance of Veillonella, Enterobacteriaceae, and Lachnospiraceae in pre-treatment fecal samples were associated with rCDI,^37^ while our results show the opposite directionality for these taxa. However, in this prior study, rCDI was determined retrospectively by review of the electronic medical record. Furthermore, all of these studies were limited by their cross-sectional study design and lack of longitudinal sampling. This is particularly limiting when considering the effects of antibiotic treatment for CDI on the gut microbiota.

There are only two published studies examining longitudinal changes in the gut microbiota in rCDI. Seekatz, *et al*. demonstrated significant differences at baseline in community structure in rCDI vs. non-rCDI patients.^17^ Notably, patients with rCDI were more likely to show greater intra-individual similarity in their gut microbiota longitudinally compared with non-rCDI patients, which is discordant with our results. However, similar to the study design by Khanna, et al., patients were retrospectively designated as having rCDI vs. recovery by chart review. Additionally, the number of and interval between samples collected was highly variable with samples collected up to 800 days after initial CDI diagnosis. Thus, our respective studies may have captured different trajectories in gut microbial changes between perturbation and reconstitution resulting in rCDI or recovery. Furthermore, our results may provide a more clinically useful model given the ability to determine risk for rCDI at baseline, or shortly after completion of antibiotics rather than weeks or months after collection of the index sample.

In the first prospective longitudinal study regarding fecal microbial changes in rCDI, Pakpour, *et al*. identified decreased Shannon diversity, depletion of Bacteroidetes and reduced abundance of *Veillonella dispar* by random forest models as being predictive of rCDI.^38^ However, the predictive capability of their model was poor (AuROC=0.68). Although there were differences in the microbial characteristics identified, this was likely related to differences in patient populations between our studies. The majority of patients enrolled by Pakpour, *et al*. identified as being of Afro-Caribbean descent and IBD patients were specifically excluded. Interestingly, these authors identified only baseline microbial characteristics as being associated with rCDI while their longitudinal samples could not discriminate between rCDI and non-rCDI. It is important to note two key methodologic differences between our respective studies. Pakpour, *et al*. collected three stool samples shortly after initiating antibiotic administration while a final sample was collected 4 days after completion of antibiotic treatment. It is possible that the gut microbiota may have been significantly disturbed by on-going antibiotic administration at this time point, which precluded discrimination between patients who recurred vs. those who recovered. Furthermore, subjects were only followed for 2-4 weeks, which is likely insufficient follow-up time to adequately determine rCDI.

In contrast, we identified distinct temporal changes in the gut microbiota that were associated with rCDI. We identified specific microbial changes 14 days after completion of antibiotics which were predictive for risk of future rCDI. Specifically, those patients whose fecal microbiota showed greater dissimilarity to their baseline samples were at highest risk for rCDI. This suggests that patients with on-going disruptions in the gut microbial ecosystem have a permissive state for recurrence of CDI. Additionally, we identified increased Ruminococcaceae and increased Enterobacteriaceae were associated with increased risk for rCDI while an increase in community richness and Faecalibacteria were protective against rCDI even after adjusting for UC status. Similar to Lachnospiraceae, Faecalibacteria are in the Clostridiales order, produce butyrate, and are associated with decreased risk for CDI.^39^ Expansion of Enterobacteriaceae, which normally occupy only a small fraction of the distal gut microbiota in healthy subjects, can be seen during periods of gut inflammation, such as in IBD or CDI.^3, 40^ Interestingly, examining the fecal microbiota at baseline or at reconstitution (14 days after completion of antibiotics) separately produced more robust models compared with longitudinal associations by generalized estimating equations, mixed models or generalized additive models in this cohort. This suggests that studying the fecal microbiota separately at baseline or after reconstitution of the microbiome, rather than the trajectory of the gut microbiota longitudinally, will be most useful in identifying those patients at highest risk for rCDI.

Prior IBD studies have identified different microbial features that are associated with worse outcomes. Using a random forest classifier, Shaw, *et al*. identified two genera, including *Coprococcus* and *Adlercreutzia*, which showed fair predictive ability to identify responders to therapy in IBD (AuROC=0.75).^41^ We identified prior hospitalization for UC in the previous year and increased Bacteroidetes at baseline as showing good predictive ability to identify patients at higher risk for subsequent UC flare in the next 180 days (AuROC=0.88). Previously, studies have demonstrated increased Bacteroidetes from mucosal biopsies of UC patients compared with controls.^42^ Some members of the Bacteroidetes phyla, such as certain *Bacteroides* species, can impair epithelial barrier function by producing mucin-degrading sulfatases or may lead to immune activation via Toll-like receptor 4.^43, 44^ It is notable that we were not able to develop a model identifying microbial characteristics longitudinally with risk for UC flare. This may reflect differences in the pathogenesis of these two different diseases, where UC may be driven more by host factors, while CDI is dependent on disruptions to the microbial ecological network. In addition, this study could have been underpowered for microbial differences in UC, and a larger cohort with increased frequency and/or longer duration of sampling could be required to develop more robust microbial-driven models of UC exacerbation given the wide variability of the IBD microbiome.^45^

It is difficult to clinically differentiate between an exacerbation of UC vs. CDI in UC patients.^46^ Our results suggest that patients with CDI have significant reductions in Shannon diversity and community richness at baseline even after controlling for UC status. Similarly, CDI patients have depletion of several OTUs at baseline, particularly in the genera Lachnospiraceae. These results suggest that baseline microbial features may be of help in differentiating between a flare and CDI in symptomatic UC patients while modulation of the microbiome in high-risk patients to increase abundance of Lachnospiraceae may be considered in future studies.

There are several notable strengths of our study, including the longitudinal study design with clinical outcome data for up to 6 months, repeated sample collection, robust demographic and clinical data, and well-defined study endpoints. However, there are limitations to our study. First, there were a small number of patients who developed UC flare, which likely limited our ability to develop a predictive model for exacerbation of UC. Secondly, we only sampled the fecal microbiota, while the mucosal-associated microbiota may provide more relevant information regarding host-microbial interactions in UC and CDI. Finally, a longer follow-up time may be required for other important adverse clinical outcomes, such as those requiring colectomy.

In conclusion, we have identified characteristic temporal changes in the gut microbiota associated with increased risk for subsequent rCDI in a cohort of patients with UC and/or CDI. Although our findings require validation, these results could have important clinical implications. The novel ability to identify patients at high risk for rCDI with over 90% accuracy using a single stool sample, either at baseline prior to initiation of antibiotics or 14 days after completion of antibiotics, could affect future clinical decision making. Inclusion of changes in microbial diversity over time adds accuracy to the model but may be less clinically useful as this requires sampling at two different time points. Clinicians could utilize this microbial-derived information to escalate preventive therapy in higher-risk patients if prospective studies validate a risk-stratified strategy of preventive therapy. Future work will focus on the mechanisms of how shifts in the gut microbiota predispose patients to rCDI, and rational design of preventive probiotics to reduce recurrence of Clostridioides difficile infection in high risk patients.

## ABBREVIATIONS

AuROC: area under the receiver operating characteristic curve
CDI: *Clostridium difficile* infection
EOA: end of antibiotics
FMT: fecal microbiota transplantation
IBD: inflammatory bowel disease
Lasso: least absolute shrinkage and selection operator
JSD: Jensen-Shannon distance
MHI: microbiome health index
OUT: operational taxonomic unit
PERMANOVA: permutational multivariate analysis of variance
rCDI: recurrent *Clostridium difficile* infection
UC: ulcerative colitis

## AUTHOR CONTRIBUTIONS

AL collected clinical data, participated in analysis and interpretation of the data, statistical analysis, and drafted the manuscript. KR participated in analyzing the data, interpreting the results, statistical analysis, and helped draft the manuscript. JL participated in acquisition of clinical data and sample processing. MG participated in acquisition of data, sample processing and data analysis. BM participated in acquisition of clinical data and sample processing. VBY participated in study design, interpreting the results, helped draft the manuscript. PDRH conceived the study, participated in subject recruitment, data acquisition, interpretation of the results, and helped draft the manuscript. All authors were involved in critical revision of the manuscript and approved the final manuscript.

## SUPPLEMENTAL METHODS

### Clinical Data Extraction

Clinical data were collected from patient interviews and manual chart review. Clinical and demographic variables included age, gender, ethnicity, current/prior smoking history, current therapy for UC, laboratory results, current/prior use of acid suppressive medications, current/prior use of probiotics, prior history of CDI, prior use of antibiotics within the past year, and hospitalization for UC in the past year.

### DNA Extraction and 16S rRNA Sequencing

All stool samples were collected and immediately stored in a −80°C freezer. Total fecal DNA was extracted using the MoBio PowerMag soil isolation kit (MoBio Laboratories). DNA libraries were prepped by the University of Michigan Host Microbiome core as previously described.^17^ Briefly, dual-index primers specific to the V4 region of the 16S rRNA gene were employed for DNA amplification.^47^ PCR mixtures contained 5 μl of primer set at 4 μM concentration, 0.15 μl Accuprime High-Fidelity *Taq*, 2 μl of 10x Accuprime PCR II buffer (Life Technologies), 11.85 μl sterile water, and 1 μl of DNA template. Cycling conditions consisted of initial denaturation at 95°C for 2 min, followed by 30 cycles of 95°C for 20 s, 55°C for 15 s, 72°C for 5 min, and then 72°C for 10 min. Samples were normalized to the lowest concentration of the pooled plates by a Sequel Prep normalization plate kit (Life Technologies). Amplicons were pooled and quantified by the Kapa Biosystems Library Quantification kit (Kapa Biosystems) while amplicon size was determined by the Agilent Bioanalyzer high-sensitivity DNA analysis kit. Libraries were then spiked with 10% PhiX to add diversity and sequenced on the Illumina MiSeq platform according to the manufacturer’s specification for 500 cycles.

### Raw Sequence Processing and Analysis

Raw sequence files were analyzed using the software package mothur (version 1.35.1).^16^ Sequences were then aligned to the V4 region using the SILVA rRNA database project (release 132).^48^ Sequences were taxonomically classified at 80% bootstrap minimum using the RDP database (release 11).^49^ Operational taxonomic units (OTUs) were clustered at 97% similarity and used for downstream community analyses. The number of OTUs (richness), Shannon index, and Jensen-Shannon distance metrics were calculated in mothur using OTU abundance. Principal coordinates analysis (PCoA) plots based on redundancy analysis were then analyzed using the R package *vegan* (version 2.5-4).^21^ Differences in community structure for each group were compared using permutational multivariate ANOVA (PERMANOVA).

## SUPPLEMENTAL TABLES

**Supplemental Table 1.**
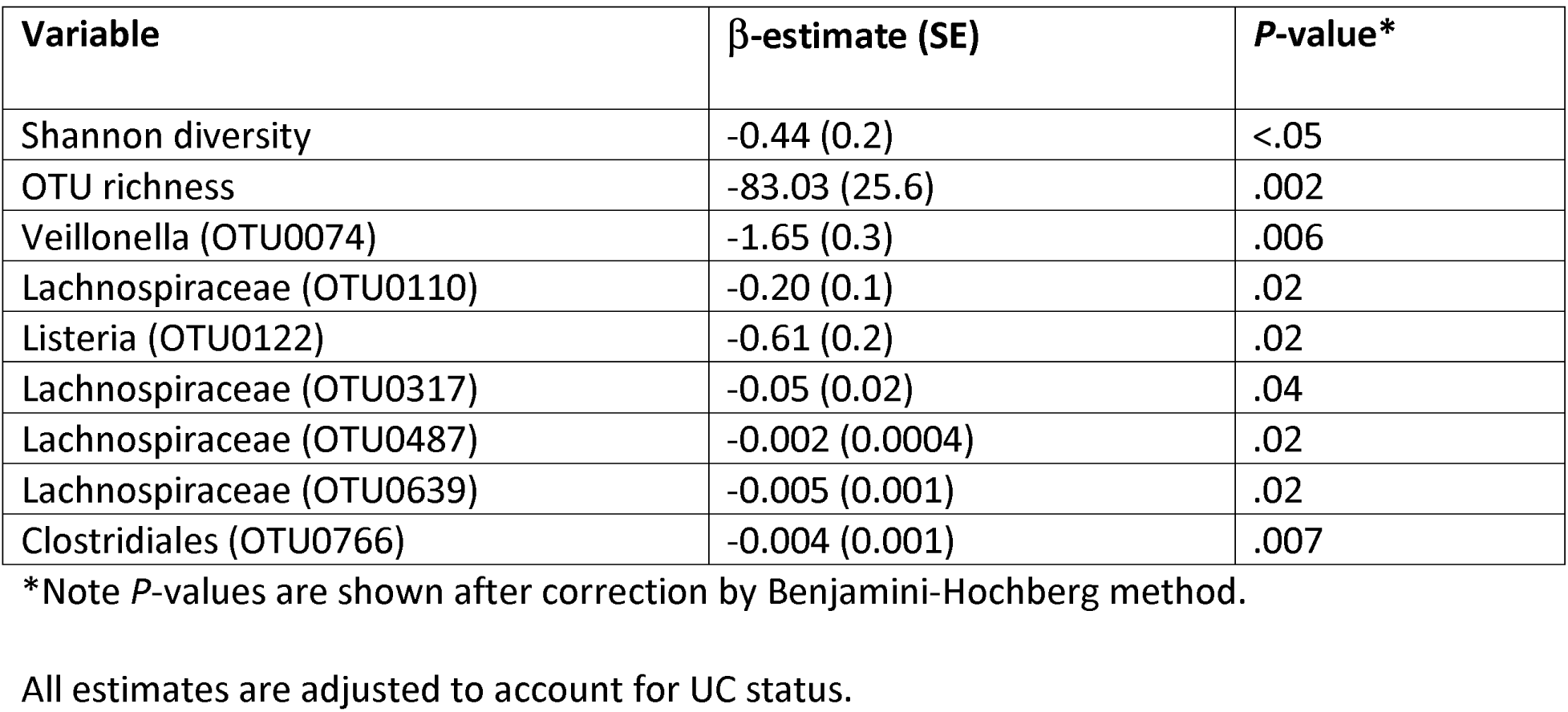
Changes in Microbial Variables at Baseline in Patients with *Clostridium difficile* Infection (CDI) Compared with Ulcerative Colitis.

| Variable | β-estimate (SE) | *P*-value* |
| --- | --- | --- |
| Shannon diversity | -0.44 (0.2) | <.05 |
| OTU richness | -83.03 (25.6) | .002 |
| Veillonella (OTU0074) | -1.65 (0.3) | .006 |
| Lachnospiraceae (OTU0110) | -0.20 (0.1) | .02 |
| Listeria (OTU0122) | -0.61 (0.2) | .02 |
| Lachnospiraceae (OTU0317) | -0.05 (0.02) | .04 |
| Lachnospiraceae (OTU0487) | -0.002 (0.0004) | .02 |
| Lachnospiraceae (OTU0639) | -0.005 (0.001) | .02 |
| Clostridiales (OTU0766) | -0.004 (0.001) | .007 |

\*Note *P*-values are shown after correction by Benjamini-Hochberg method.
All estimates are adjusted to account for UC status.

**Supplemental Table 2.**
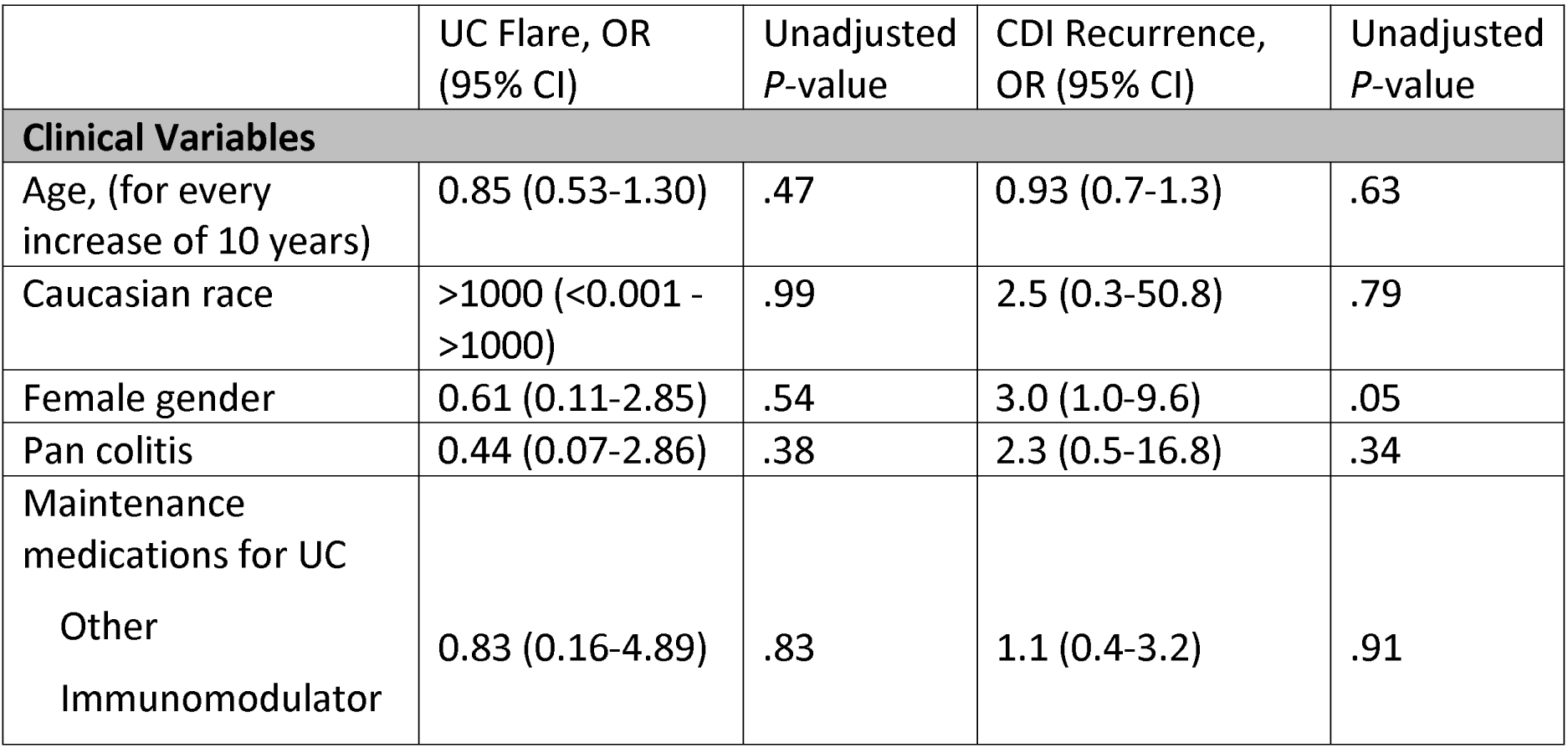
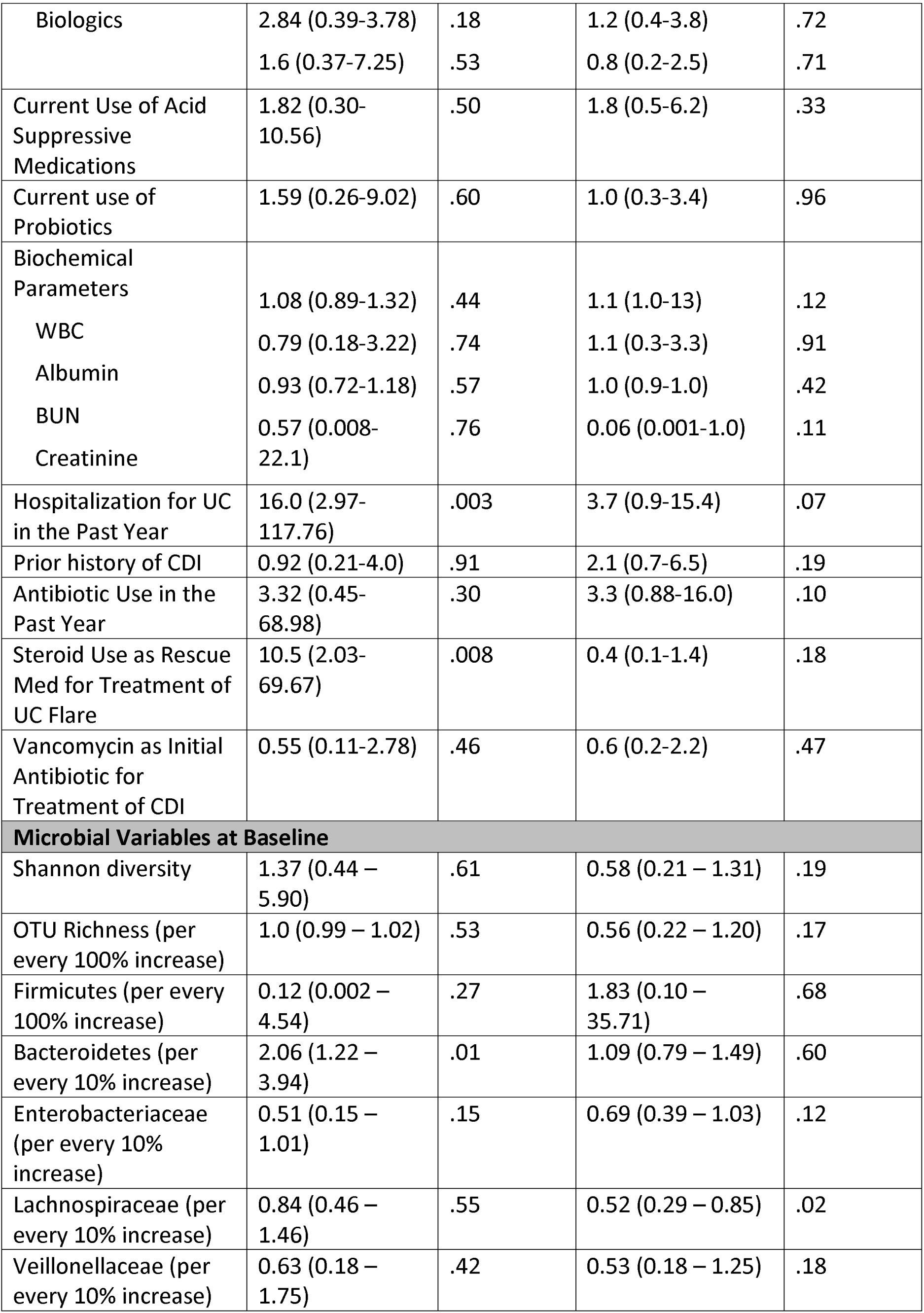
Unadjusted Odds Ratios for Baseline Clinical and Microbial Variables Associated with Recurrent *Clostridium difficile* Infection (rCDI) and Ulcerative Colitis (UC) Flare.

**Supplemental Table 3.** Lasso Regression Model of Clinical and Microbial Variables Associated with Risk for Recurrent *Clostridium difficile* Infection (rCDI) at Baseline.

| | β-Estimate |
| --- | --- |
| ***Clinical and Microbial Variables Associated with Increased Risk for rCDI*** |  |
| Hospitalization for UC in prior year | 1.72 |
| Female gender | 1.26 |
| Verrucomicrobia | 0.07 |
| Ruminococcaceae | 0.06 |
| ***Microbial Variables Protective Against rCDI*** |  |
| Eubacteriaceae | -12.32 |
| Lachnospiraceae | -0.07 |
| Veillonellaceae | -0.06 |
| Enterobacteriaceae | -0.04 |

$\beta$ -estimates for all microbial variables are represented per every 1% increase in relative abundance.

**Supplemental Table 4.**
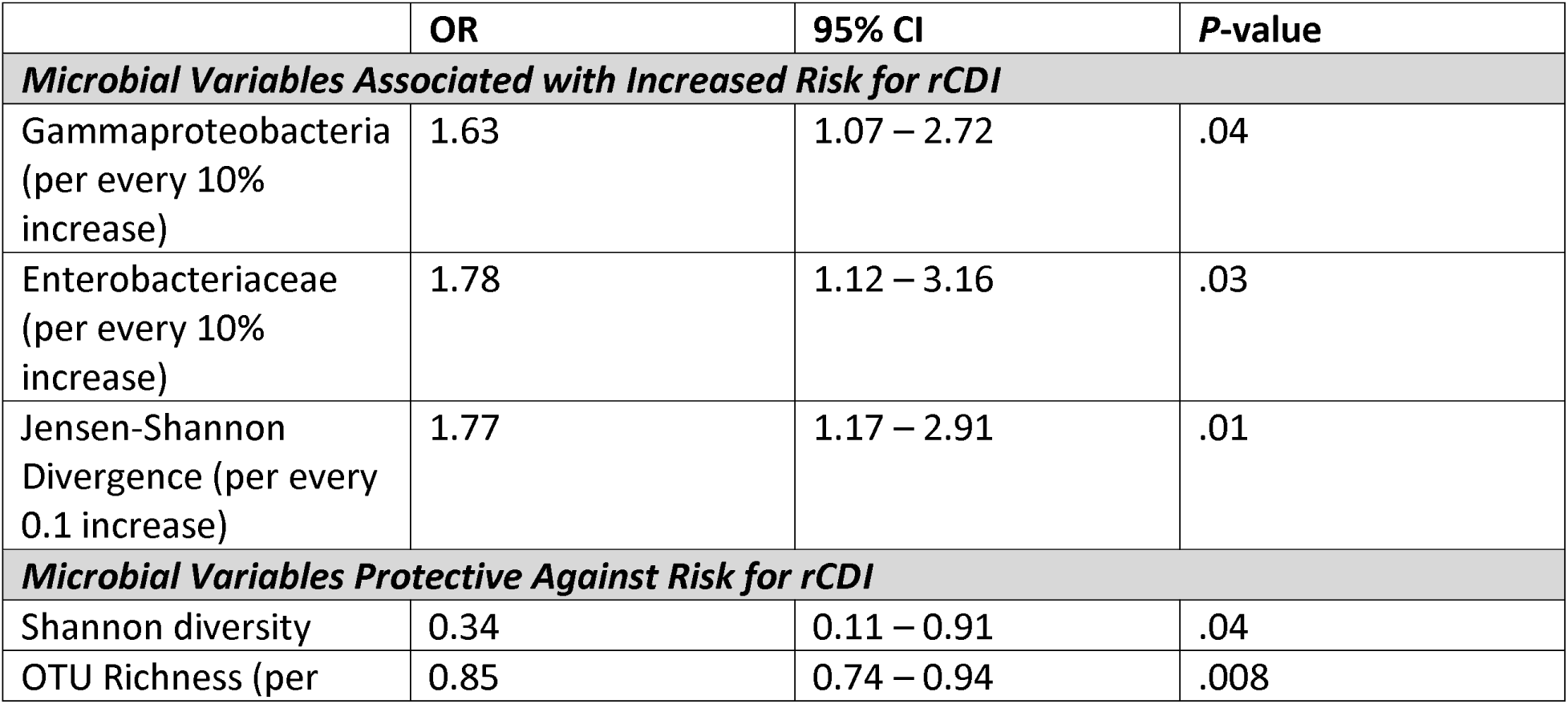
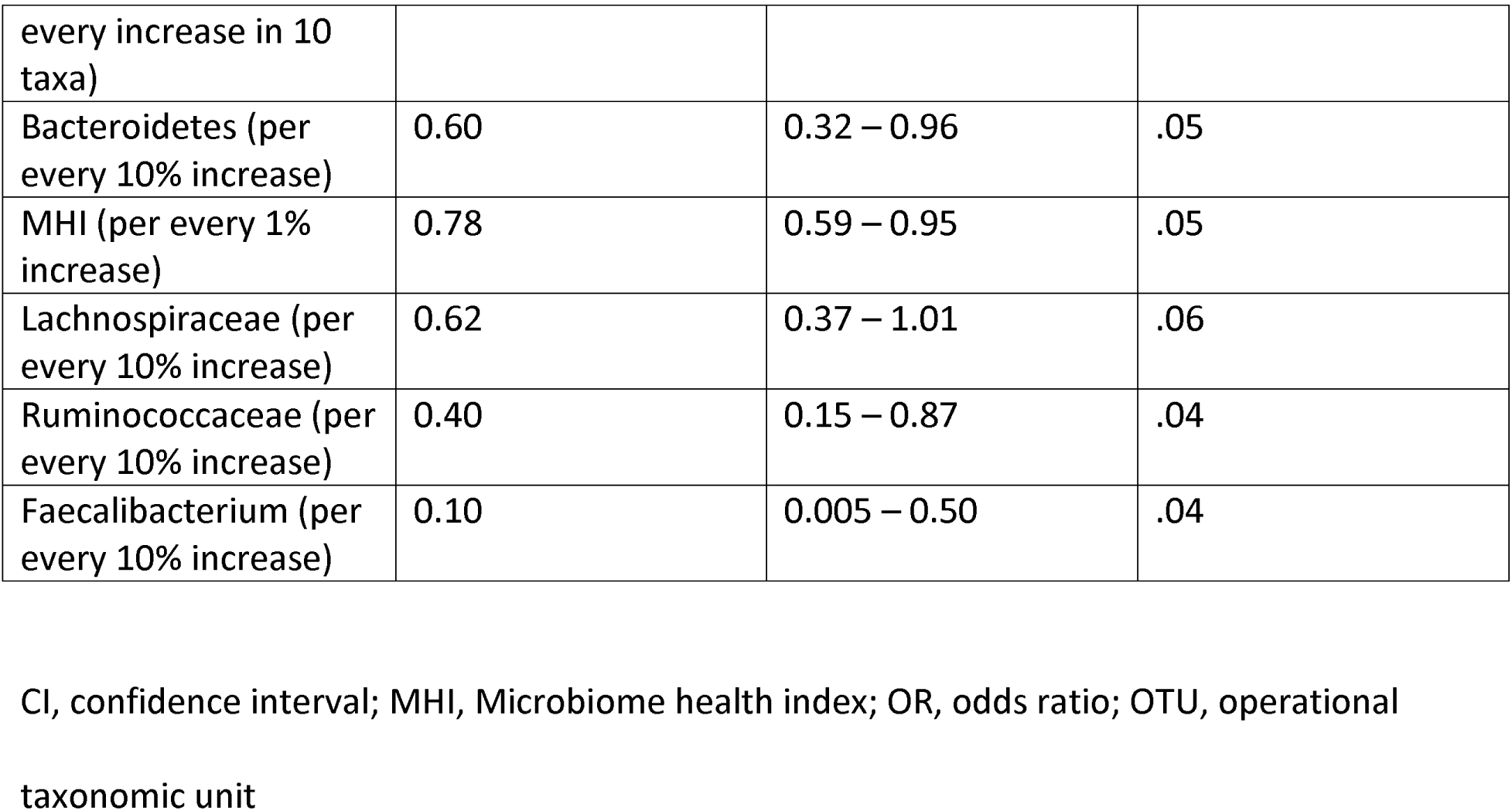
Unadjusted Odds Ratios for Microbial Variables Associated with Recurrent *Clostridium difficile* Infection (rCDI) 14 days After Completion of Antibiotics.

**Supplemental Table 5.** Lasso Regression Model of Clinical and Microbial Variables Associated with Risk for Recurrent *Clostridium difficile* Infection (rCDI) 14 Days After Completion of Antibiotics.

**Antibiotics**
| | β-Estimate |
| --- | --- |
| ***Clinical and Microbial Variables Associated with Increased Risk for rCDI*** |  |
| Jensen Shannon Diversity (relative to baseline sample) | 5.84 |
| Female gender | 1.53 |
| Shannon diversity | 1.39 |
| Ruminococcaceae | 0.20 |
| Enterobacteriaceae | 0.04 |
| ***Microbial Variables Protective Against rCDI*** |  |
| Faecalibacterium | -0.29 |
| OTU richness | -0.02 |

$\beta$ -estimates for OTU richness is represented for every increase in 1 taxa while microbial
variables are represented per every 1% increase in relative abundance.

## SUPPLEMENTAL FIGURES

**Supplemental Figure 1.** Consort Flow Diagram.

**Supplemental Figure 2. Study Design and Fecal Sample Collection Timeline.**

Subjects with ulcerative colitis (UC) and *Clostridium difficile* infection (CDI) as well as non-inflammatory bowel disease (IBD) controls with CDI provided stool sample 1 (SS1) at day 0 when they had positive CDI testing but prior to initiation of antibiotics. Additional samples were provided at the end of antibiotic therapy (SS2) and at the end of antibiotic therapy plus 14 days (SS3). Subjects were then contacted at 60, 120, and 180 days to assess for recurrent UC flare and/or recurrent CDI. A third cohort of patients with UC flare without CDI provided stool samples at baseline (SS1) and again at day 30 (SS2) when their disease was in remission.

**Supplemental Figure 3. Baseline Microbial Variables Associated with Ulcerative Colitis (UC) Flare.**

Increased relative abundance of Bacteroidetes (presented on a logarithmic scale) (OR=2.06 for every 10% increase, *P*=.01) was significantly associated with risk for ulcerative colitis (UC) flare. Data are shown as violin plot with accompanying boxplot and whiskers indicating median, interquartile range (IQR) and 1.5 x IQR of the median.

**Supplemental Figure 4. Patients with Recurrent *Clostridium difficile* Infection (rCDI) and Ulcerative Colitis (UC) Flare Demonstrate Non-Linear Dynamic Changes in the Fecal Microbiota.**

Longitudinal modeling of the fecal microbiota demonstrated a non-linear pattern in this patient cohort. Representative examples include (A) Shannon diversity for patients with and without recurrent *Clostridium difficile* infection (rCDI) as well as (B) patients with and without ulcerative colitis (UC) flare are shown at baseline, end of antibiotics (EOA) and end of antibiotics plus 14 days (EOA+14d). Individual subjects are represented by different colored lines. (C) Operational taxonomic unit (OTU) richness in patients with and without rCDI as well as (D) in patients with and without UC flare is also shown longitudinally. A locally estimated scatterplot smoothing (LOESS) curve has been fitted to the data and depicted by the black line with error bars in gray.

