## Supplemental Figure 1 for "Temporal Gut Microbial Changes Predict Recurrent *Clostridium difficile* in Patients with and without Ulcerative Colitis"

**CONSORT Flow Diagram**

**Allocation**

**Analysis**

**Follow-Up**

**Enrollment**

Assessed for eligibility (n=208)

Excluded (n=121)

♦  Not meeting inclusion criteria (n=3)

♦  Declined to participate (n=118)

Reasons: Did not provide all three stool samples (n=30)

Lost to follow-up (give reasons) (n=30)

Discontinued intervention (n=0)

Enrolled (n=87)

♦ Did not enroll (n=0)

Analysed (n=57)
♦ Excluded from analysis (n=0)
