## Supplementary figures and images for "Temporal Gut Microbial Changes Predict Recurrent *Clostridium difficile* in Patients with and without Ulcerative Colitis"

### Supplemental Figure 2

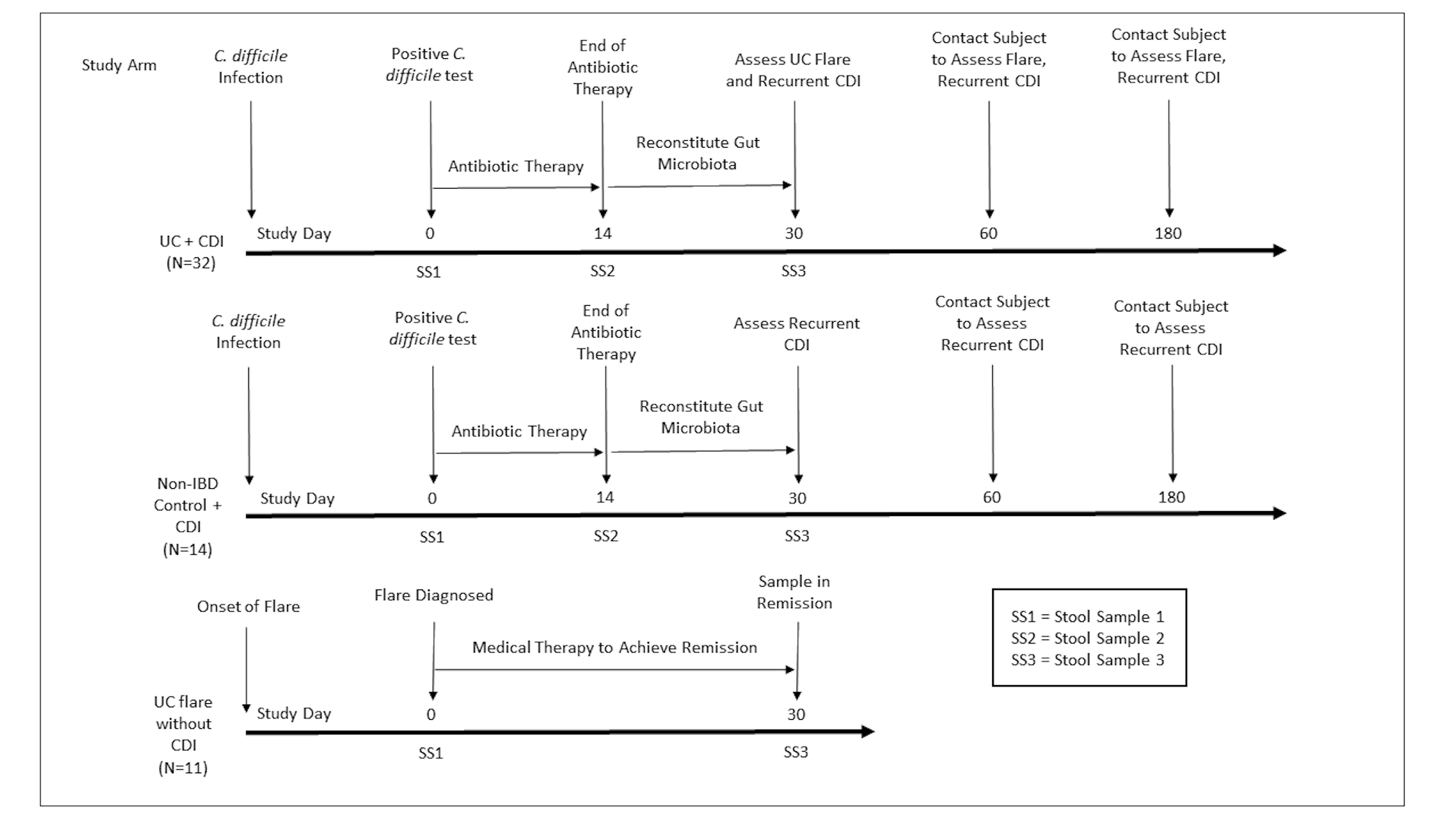

### Supplemental Figure 3

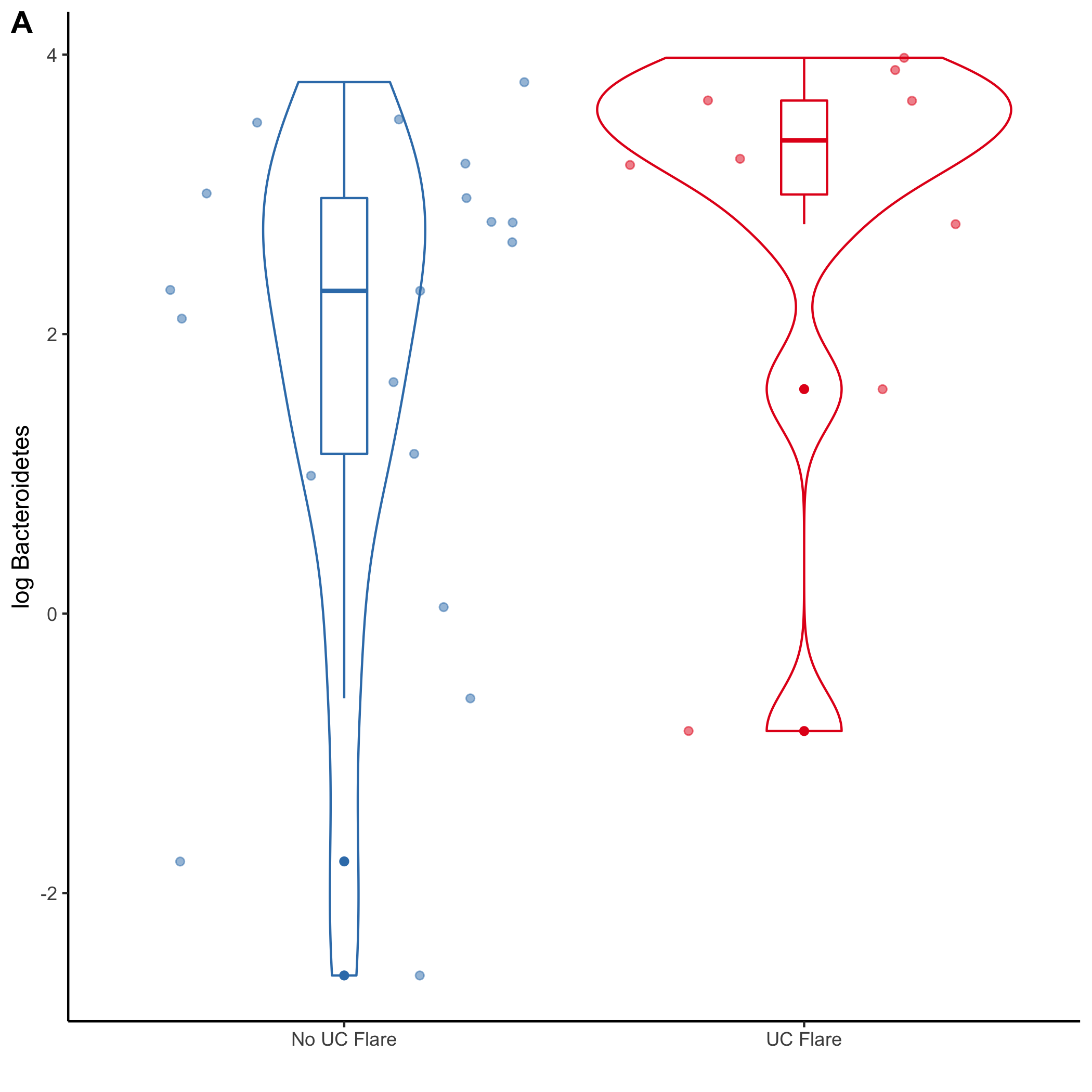

### Supplemental Figure 4

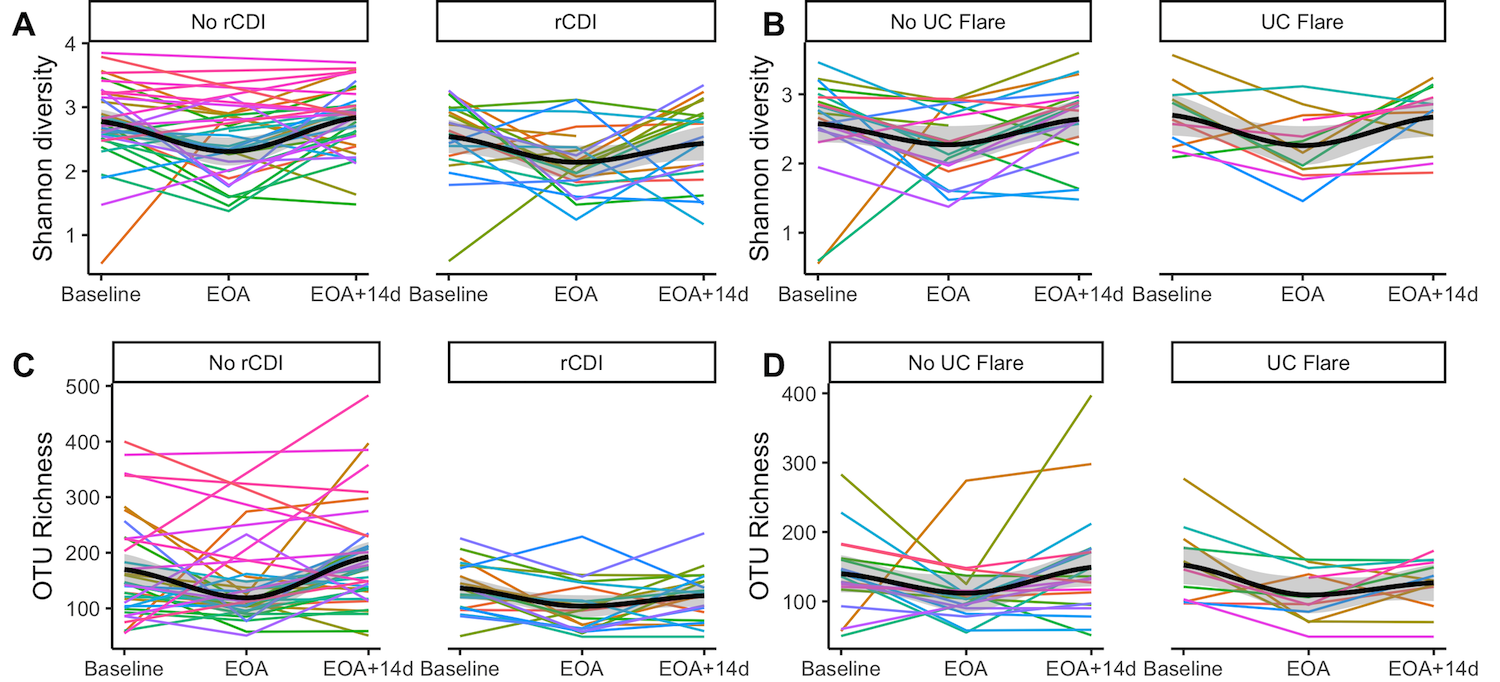
